# Acute effects of psilocybin on the dynamics of gaze fixations during visual aesthetic perception

**DOI:** 10.1101/2023.10.27.564413

**Authors:** Stephanie Muller, Federico Cavanna, Laura de la Fuente, Nicolás Bruno, Tomás Ariel D’Amelio, Carla Pallavicini, Enzo Tagliazucchi

## Abstract

**Rationale:** Serotonergic psychedelics are remarkable for their capacity to induce variable yet reproducible modifications to human consciousness The most salient acute effects of these compounds include perceptual alterations, predominantly in the visual domain, yet to date these alterations have been mostly documented only by subjective reports.

**Objectives:** We used eye-tracking to quantify the effects of low vs. high doses of psilocybin mushrooms on the eye movements that underlie the exploration of complex visual stimuli, focusing on the particular case of aesthetic perception.

**Results:** Following a double-blind placebo-controlled design under semi-naturalistic conditions, we demonstrated that high doses of psilocybin result in a more local visual exploration of paintings, and thus in a less entropic fixation probability distribution. Participants reported heightened emotional response and state of flow under the high dose condition.

**Conclusions:** These findings are consistent with an effect of psilocybin on gaze fixation mediated by altered perception of low-level visual information, such as textures, shapes and colors. Our work also highlights the possibility of investigating psychedelics by addressing their effect on behavior under complex naturalistic conditions, which contributes to maintaining subject engagement while also increasing the ecological validity of the findings.

## Introduction

Moderate to high doses of classic psychedelic substances (i.e. LSD, psilocybin, DMT and mescaline) are known to produce profound alterations in consciousness, including acute changes in perception, emotion, cognition, and the sense of self (Nichols, 2016; Preller & Vollenweider, 2018; van Elk & Yaden, 2022). The effects of serotonergic psychedelics on human subjective experience show consistency despite individual variation, drug dose and non-pharmacological inter-individual factors (Studerus et al., 2012). Perceptual alterations are among the most prominent and commonly reported acute effects induced by psychedelics. These alterations encompass visual and auditory distortions, changes in body schema and tactile perception, the onset or enhancement of synesthesia and disrupted perception of time (Nichols, 2016; Preller & Vollenweider, 2018; van Elk & Yaden, 2022). Changes in perceptual meaning are also frequently reported, with some items in the environment becoming more salient and personally relevant than usual (Kometer & Vollenweider, 2018; Liechti et al., 2017; Studerus et al., 2010).

Early research attempted to provide insight into how psychedelics affect perception and cognitive functions (e.g., memory, attention, decision-making, etc.) (Nichols & Walter, 2020). However, several of these early studies present limitations such as unrefined methodological standards, small sample sizes, inconsistent blinding procedures and control groups, non-standardized dosages and heterogeneous rating scales (Aday et al., 2021; van Elk & Fried, 2023). In addition, the tools used to study the effects of psychedelics in the human brain were limited by the technology available at the time. Researchers relied mainly on subjective self-reports from participants, observational data and animal models (de Deus et al., 2023); however, as psychedelic research declined in the 70s and 80s, new tools and technologies emerged. The development of neuroimaging techniques such as MRI and fMRI allowed researchers to visualize and study brain structure and function in vivo, while eye tracking technology improved, becoming more widely available and affordable, and allowing accurate measurements of gaze direction capable of informing cognitive processes underlying perception and attention (Borys & Plechawska-Wójcik, 2017). Even though recent studies have begun to re-examine the psychophysical effects of psychedelics and their capacity to alter perception, most of this research attempts to correlate brain imaging data with self-reported sensory alterations (Dos Santos et al., 2016; de Deus et al., 2023). Thus, a gap in knowledge remains concerning the impact of psychedelic drugs on objective behavioral metrics, such as the dynamics of sensory engagement with the environment.

In this double-blind placebo-controlled study conducted in a semi-natural setting, we used eye tracking to measure the dynamics of eye fixations as participants freely explored a selection of classical paintings while under the acute effects of high and low doses of psilocybin present in *Psilocybe cubensis* mushrooms. Additionally, we recorded self-reported evaluations of emotional valence and beauty ratings for each painting. As a first attempt to investigate the effects of psychedelics on the dynamics of visual perception, our study is inherently exploratory. However, our methodology was deliberately chosen to foster a high level of engagement while minimizing possible sources of stress and distraction (Bălăeţ, 2022). Moreover, the choice of classical paintings was based on scientific evidence and anecdotal reports indicating that psychedelics positively interact with aesthetic experience (McGlothlin et al., 1967; Spee et al., 2018; van Elk et al., 2022; Aday et al., 2023). Subjects participated in this experiment during their natural use of the drug, with the agreement to engage in a control condition which was prepared and randomized by a third party. Because of this semi-naturalistic design, participants benefited from a familiar and comfortable research setting, which could attenuate their overall distraction and contribute to mitigate confounding factors related to stress and anxiety, potentially amplified by the acute effects of the drug (Studerus et al., 2012).

To complement the exploratory nature of this study, we adopted a hypothesis based on the Relaxed Brain Under Psychedelics (REBUS) model, a theoretical proposal of how psychedelics alter brain function to elicit changes in consciousness and cognition. According to this model, psychedelics bring about the revision of heavily weighted high-level priors (Carhart-Harris and Friston 2019). During the visual perception, this could be manifest as a reduced focus on salient features (i.e. informative priors for image recognition and categorization), and thus as a more unbiased and homogeneous sampling of the visual stimuli – or equivalently, as an increased entropy in the spatial distribution of gaze fixations (Bartels and Zeki 2004; Hasson et al. 2004; Fischera et al. 2013). These predictions can be tested by comparing the probability distribution of fixations between the high and low dose conditions.

## Materials and methods

### Participants

Twenty-three participants (four females, 31 ± 4 years, 72 ± 15 kg [mean ± STD]) were recruited through word of mouth and social media advertising. To participate, individuals were instructed to contact the provided number via WhatsApp, and subsequently received a phone call from the researchers. During the call, the researchers provided a brief explanation of the details and purpose of the experiment, as well as the inclusion and exclusion criteria. Participants were then given a full written explanation of the study and a copy of the informed consent form. To determine eligibility, all participants underwent a psychiatric interview to screen for exclusion criteria, which are detailed in the supplementary material and summarized below. After this screening, participants and researchers agreed on a date for the start of the experiment. Participants reported having 15 ± 13 prior experiences with serotonergic psychedelics, of which 3.1 ± 2.4 were considered challenging [mean ± STD]. All participants had normal or corrected-to-normal vision.

This study was conducted in accordance with the Declaration of Helsinki and approved by the Research Ethics Committee at the Universidad Abierta Interamericana (Buenos Aires, Argentina), protocol number 0-1068. All participants gave written informed consent and received no financial compensation for their participation in the experiment.

### Inclusion and exclusion criteria

Participants were required to have at least two prior experiences with a dose equal to or exceeding 3 g of dried psilocybin mushrooms. To participate in this research protocol, subjects volunteered to partake in a series of tests under the effects of psilocybin mushrooms and in the presence of four members of the research team. Subjects who consumed serotonergic psychedelics during the 15 days prior to the dosing day were not included in the study. The same applied for all psychoactive substances (including alcohol, caffeine and tobacco) for a period of 24 hours prior to the dosing day.

A non-diagnostic psychiatric interview was conducted according to the guidelines by Johnson et al. (2008). Subjects who fulfilled DSM-5 criteria for the following disorders were excluded from the experiment: schizophrenia or other psychotic disorders, type 1 or 2 bipolar disorder (including first- and second-degree relatives), substance abuse or dependence in the past 5 years (excluding nicotine), depressive disorders, recurrent depressive episodes, obsessive-compulsive disorder, generalized anxiety disorder, dysthymia, panic disorder, bulimia or anorexia, as well as subjects with a history of neurological disorders. Pregnant women and subjects under psychiatric medication of any kind were excluded, as were subjects with dysfunctional states as measured by the DASS scale (Lovibond and Lovibond 1995).

### Experimental design and setting

This study followed a randomized, double-blind placebo-controlled, within-subject design. The experiment was divided into two parts, one corresponding to the dosing condition (3 g of ground and homogenized dried psilocybin mushrooms in gel capsules) and one corresponding to the active placebo condition (0.5 g of ground and homogenized dried psilocybin mushrooms mixed with 2.5 g of an edible mushroom in gel capsules). Conditions were separated by an interval of one month to attenuate potential tolerance effects, and were randomized by a third party to maintain blinding, ensuring that neither the subjects nor the researchers knew the identity of the capsules (Cavanna et al., 2022). The experiment took place in the comfortable setting of a house and in the presence of the participant and the team of researchers. Two participants opted to stop the eye tracking task during the high dose condition and were therefore excluded from the following analysis; however, no adverse effects were reported.

### Acute effects

During the acute effects, subjects used a visual analogue scale (VAS) to provide their subjective intensity report for the following items: “Sounds influence what I see”, “My sense of size and space is distorted”, “I feel unusual bodily sensations”, “I see geometric patterns”, “Edges seem warped”, “I see movement in things that aren’t really moving”, “Things look strange”. The VAS used in this experiment is the abbreviated form of a previously used version (Pallavicini et al. 2021; Cavanna et al. 2022). which is restricted to include items that focus on the effects of the drug on perception. The VAS was completed four times, the first one hour after dose ingestion and then every one hour until the effects of the drug subsided.

### Self-reported scales and questionnaires

Two days prior to the dosing day of each condition, subjects completed the following self-reported scales designed to assess different psychological traits: State-Trait Anxiety Inventory (STAI-Trait) (Spielberger, 1983), Big Five Inventory (BFI) (John et al., 1991), Tellegen Absorption Scale (TAS) (Tellegen & Atkinson, 1981) and Short Suggestibility Scale (SSS) (Kotov et al., 2004). A battery of self-reported scales was administered both before dosing and immediately after the acute effects, including the State-Trait Anxiety Inventory (STAI-State) (Spielberger, 1983), the Positive and Negative Affect Schedule (PANAS) (Sandin et al., 1999), and the Psychological Well-being Scale (BIEPS) (Luna et al., 2020). Additionally, before dosing, subjects completed scales assessing expectations (EXP) and contextual factors prior to psychedelic experiences (PRE) (Haijen et al., 2018), and after the acute effects they completed the Mystical Experience Scale (MEQ) (MacLean et al., 2012) and the Altered States of Consciousness Scale (5D-ASC) (Studerus et al., 2010). Participants also completed the Aesthetic Experience Questionnaire (AEQ) (Wanzer et al., 2020) on the dosing day immediately after completing the eye-tracking task, which is the primary questionnaire that is reported in this work. See the supplementary material for a detailed description of the other questionnaires..

### Stimuli

A total of 30 paintings were selected from a collection of 100 paintings published by the BBC (https://www.bbc.co.uk/programmes/p009ypzq), ranging from 12th century China to the 50s. This selection was performed to represent multiple styles, themes and artistic periods, and to avoid particularly popular or famous works that are likely to be well-known by the participants. The list of the authors and titles of the selected paintings can be found in the supplementary material. Images had a maximum resolution of 1280 × 768 pixels and were presented in front of an homogeneous gray background.

### Eye tracking

The measurements reported below were conducted as part of a larger experiment designed to investigate the effects of psychedelics on perception, creativity, language and music production, as well as the associated electrophysiological correlates measured with EEG, which will be presented in a series of future reports.

For the eye tracking task, participants were seated 60 cm from a centrally located monitor with their heads supported by a custom made chin rest to prevent head movement. The tracker was placed between the subject and the screen. The monitor had a resolution of 1920 × 1080 pixels and a size of 31 × 17.5 cm. While presenting the visual stimuli, the spatial and temporal coordinates of the gaze location were recorded using the video-based Gazepoint GP3 HD eye tracker, with a temporal resolution of 150 Hz and a visual angle accuracy of 0.5 - 1.0 degrees. Both the eye tracker control and the presentation of the different stimuli were performed with a custom script written in Python using the open-source library PsychoPy (https://www.psychopy.org/).

The experiment started one hour after the participants consumed the capsules that contained either the low dose or the high dose of active mushroom material. The outline of the experiment is presented in Figure 1. Each block (T) began with the presentation of a fixation cross for 2 s; next, the stimulus was presented for 30 s. Two self-reported VAS were then presented to assess the subject’s emotional valence and beauty rating of the previously presented painting. This process was repeated until all stimuli were shown (30 blocks). To avoid changes in the measurement accuracy over time, a standard eye tracker 5-point calibration procedure was performed at the end of each fifth block.

**Figure 1.**
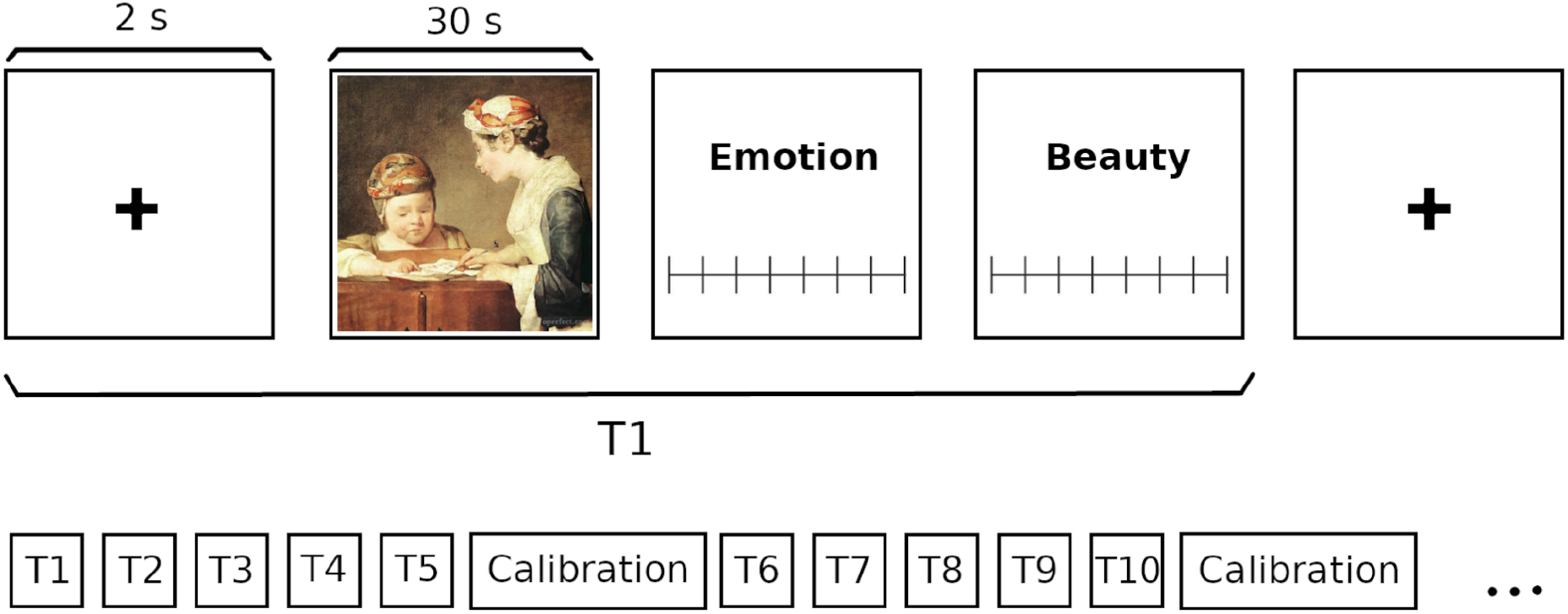
Outline of the eye tracking task. After the initial calibration, the first block (T1) begins. Each block consists of the following steps: a fixation cross is displayed for 2 s, followed by the presentation of the stimulus for 30 s, during which participants freely explore the painting while their gaze is being recorded by the eye tracker. Emotional valence and beauty ratings were assessed after each painting. This procedure was repeated for the 30 paintings, with intermediate eye tracker calibration every 5 blocks.

### Statistical fixation metrics

Due to insufficient data quality (predominantly coordinates outside those corresponding to the screen, suggestive of lack of engagement with the task) the data of 6 participants were excluded from the analysis presented in this and the following section.

After recording the coordinates of the gaze position, fixations were detected automatically using a velocity-based algorithm (Engbert and Kliegl 2003) with minimum durations of 150 ms (Manor and Gordon 2003). This resulted in the average horizontal and vertical positions (*x*_*i*_, *y*_*i*_), as well as the time of the i-th fixation (*t*_*i*_), with the positions recorded in pixels and the time in seconds. Since the dimensions of the selected paintings varied, all fixations were standardized to a common dimension of 1280 × 768 pixels to allow straightforward comparison of distances. Fixations outside the images were removed. For each image, participant and condition, the following measures were calculated: the total number of fixations within the image (N), the mean distance between fixations (ds), the mean time between fixations (dt) and the mean distance between all pairs of fixations (ds^2^). These metrics are outlined in Figure 4A, and are computed as follows:

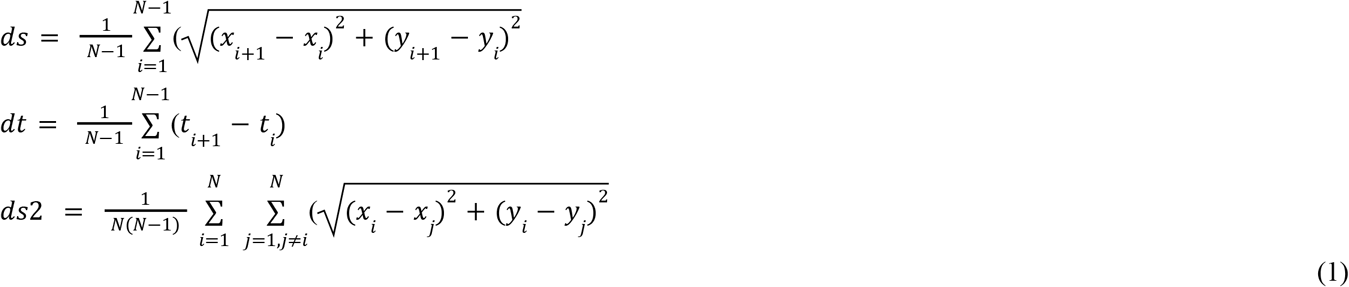

Additionally, the standard deviation was also calculated for each image for the distances between consecutive fixations *σ*(*ds*)) and the distances between all pairs of fixations (*σ*(*ds*^2^)), as follows:

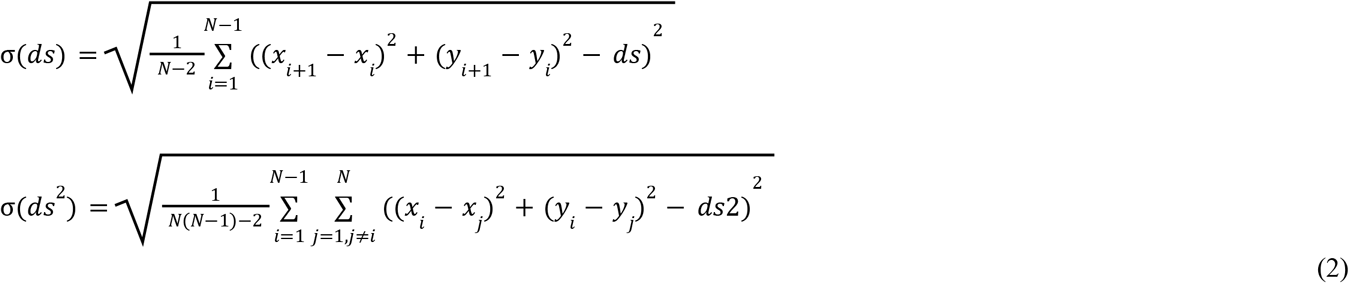

These measures were then averaged across images, resulting in a mean value for each participant and dosing condition.

### Shannon Entropy

The Shannon entropy *SE*) of the 2D fixation probability distribution was calculated for each participant and dosing condition, according to the formula by Shannon (1948):

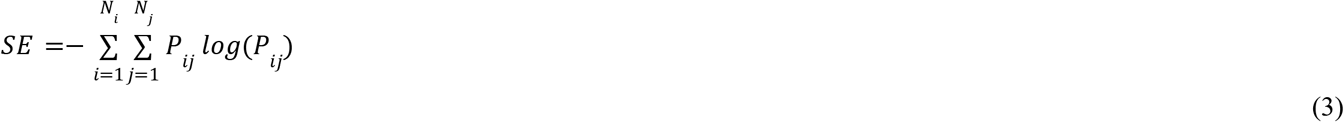

Where *N*_*i*_ and *N*_*j*_ are the total number of bins in the x and y dimensions, respectively, and *p*_*ij*_ is the value of the fixation probability distribution corresponding to the bin at position *i,j*, calculated as the number of fixations inside that bin divided by the total number of fixations.

### Statistical analyses

The results of the questionnaires from both dosing conditions were compared using Student’s t-test for paired samples, as implemented in Python’s *scipy* library (https://scipy.org). Frequentist methods were complemented using Bayesian statistics to compare the evidence in favor of the null hypothesis with that in favor of the alternative hypothesis. We computed the Bayesian statistic BF10 (Bayes factor in favor of the alternative hypothesis over the null hypothesis) as implemented in Python’s *pingouin* library (https://pingouin-stats.org) (Held & Ott, 2018). Eye-tracking measurements and VAS scores from both dosing conditions were compared using non-parametric paired Wilcoxon signed-rank tests, using Python’s *scipy* library. For a measure of the effect size, we computed the matched pairs rank-biserial correlation (RBC) as implemented in Python’s *pingouin* library. In the figures, all boxplots extend from the lower to upper quartile values with a line at the median; the whiskers extend from the upper/lower quartiles up to 1.5 times the interquartile range. Single points scattered over the boxplots represent data points from individual subjects.

## Results

Participants correctly unblinded the experimental condition in 42 of the 46 measurement days (unblinding rate of 91%), with the same percentage for high dose and active placebo condition. The results of all the self-reported scales and questionnaires (except the AEQ, shown below) appear in the supplementary material (table S1). As expected, the high dose condition resulted in significant increases across all items of the 5D-ASC and MEQ30 questionnaires. Moreover, the majority of the negative results also presented BF10 factors below 1/3, interpretable as evidence in favor of the alternative hypothesis (i.e. no effect of the psilocybin).

### Acute effects

To obtain an index representing the overall intensity of the acute effects experienced by subjects, we computed the sum of all the VAS items at each measurement time point. As expected, all four VAS total scores were significantly higher for the high dose of psilocybin vs. the active placebo (low dose). Figure 2 shows the temporal evolution of the total VAS score, as well as the scores for each individual item, averaged across participants for both dosing conditions. For all VAS scores, a peak in the intensity of the effects appeared between 2 to 3 hours after drug intake, with a tendency to decrease 4 hours. This figure also indicates the time interval when the eye tracking task was performed, as well as the time point when the AEQ was completed by the participants.

**Figure 2.**
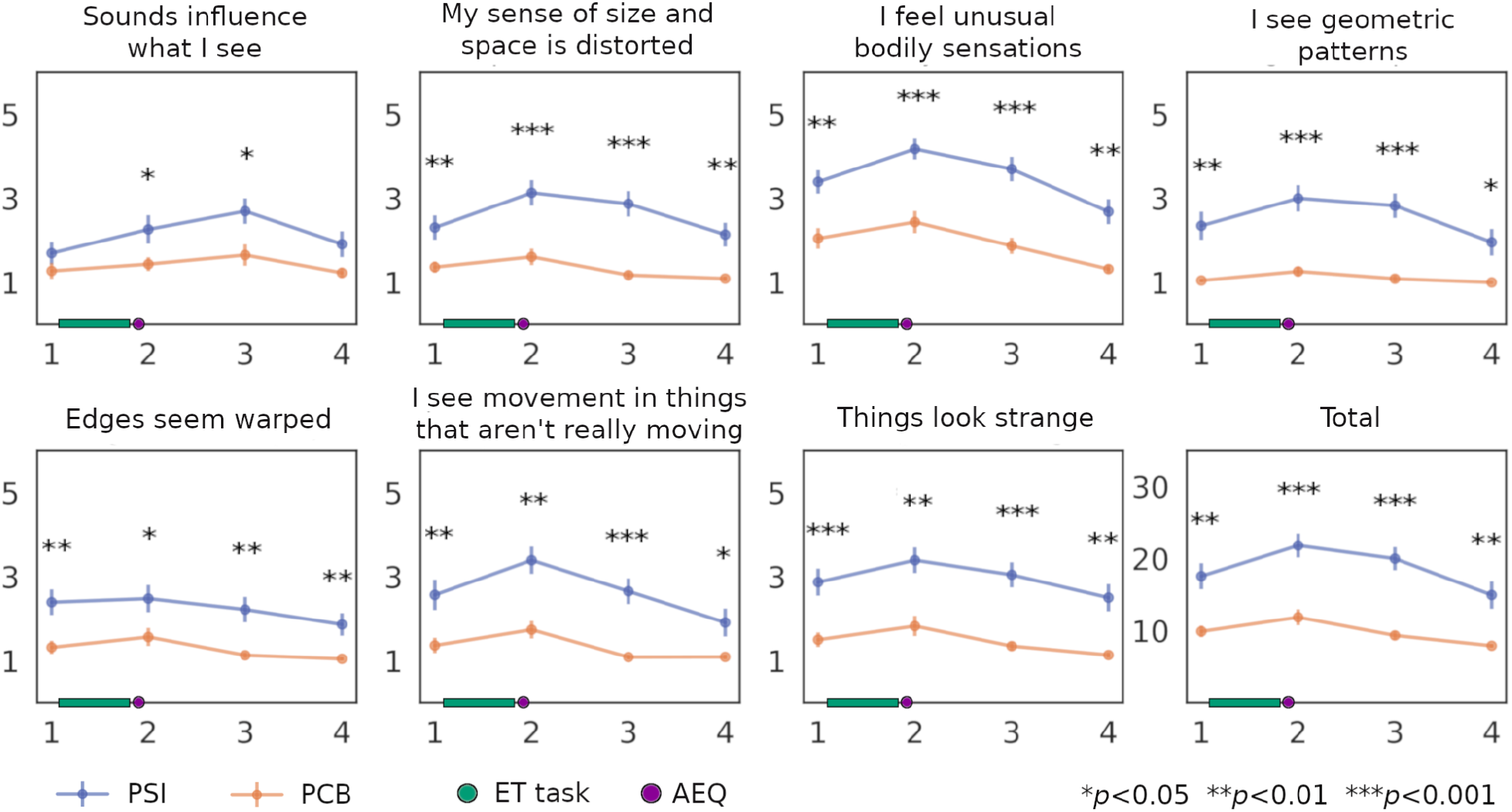
Acute effects measured using individual VAS items and the overall intensity of the experience given by the sum of all the items. Results are shown for each measurement time point, with consecutive measurements separated by one hour. The points indicate the mean across participants and the vertical lines the standard error of the mean (SEM). The time during which the eye tracking task (ET) took place and when the AEQ was completed are both indicated in the x-axis. Statistical significance is indicated using asterisks (Wilcoxon signed-rank tests).

### Subjective ratings during the perception task

The scores for the six factors of the AEQ were compared between the dosing conditions using Student t-tests, revealing that the emotional response factor and the flow of experience factor scored significantly higher in the high dose condition (Figure 3a). The emotional valence and perceived beauty VAS scores were averaged across the images for each participant and compared using Wilcoxon signed-rank tests, showing no significant differences between dosing conditions (Figure 3b).

**Figure 3.**
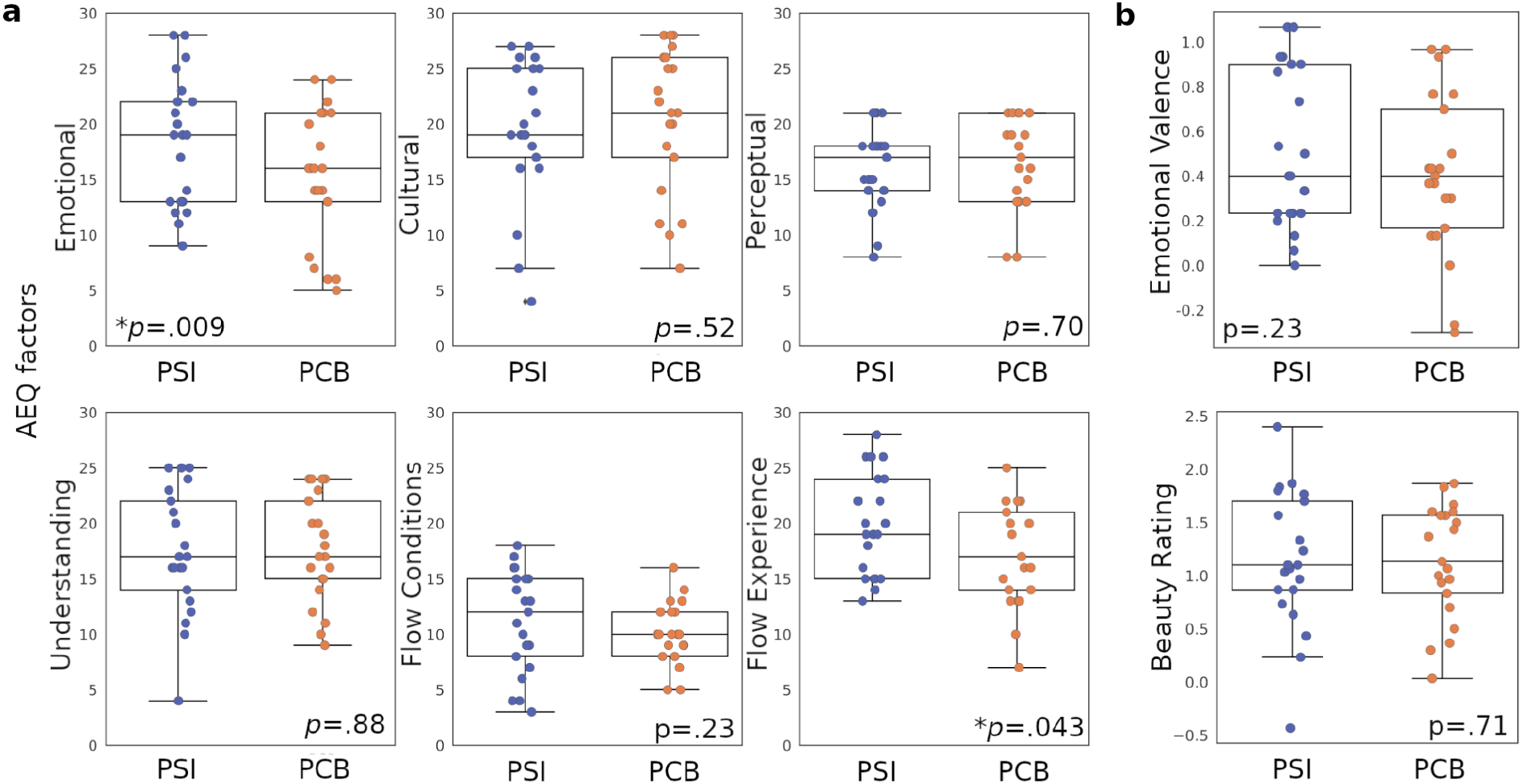
Results of the subjective reports obtained during the aesthetic perception task. **a)** Mean AEQ scores for each sub-scale, averaged across stimuli. **b)** Emotional valence and perceived beauty VAS scores, averaged across stimuli. The p-values computed using Wilcoxon signed-rank tests are shown as insets.

**Figure 4.**
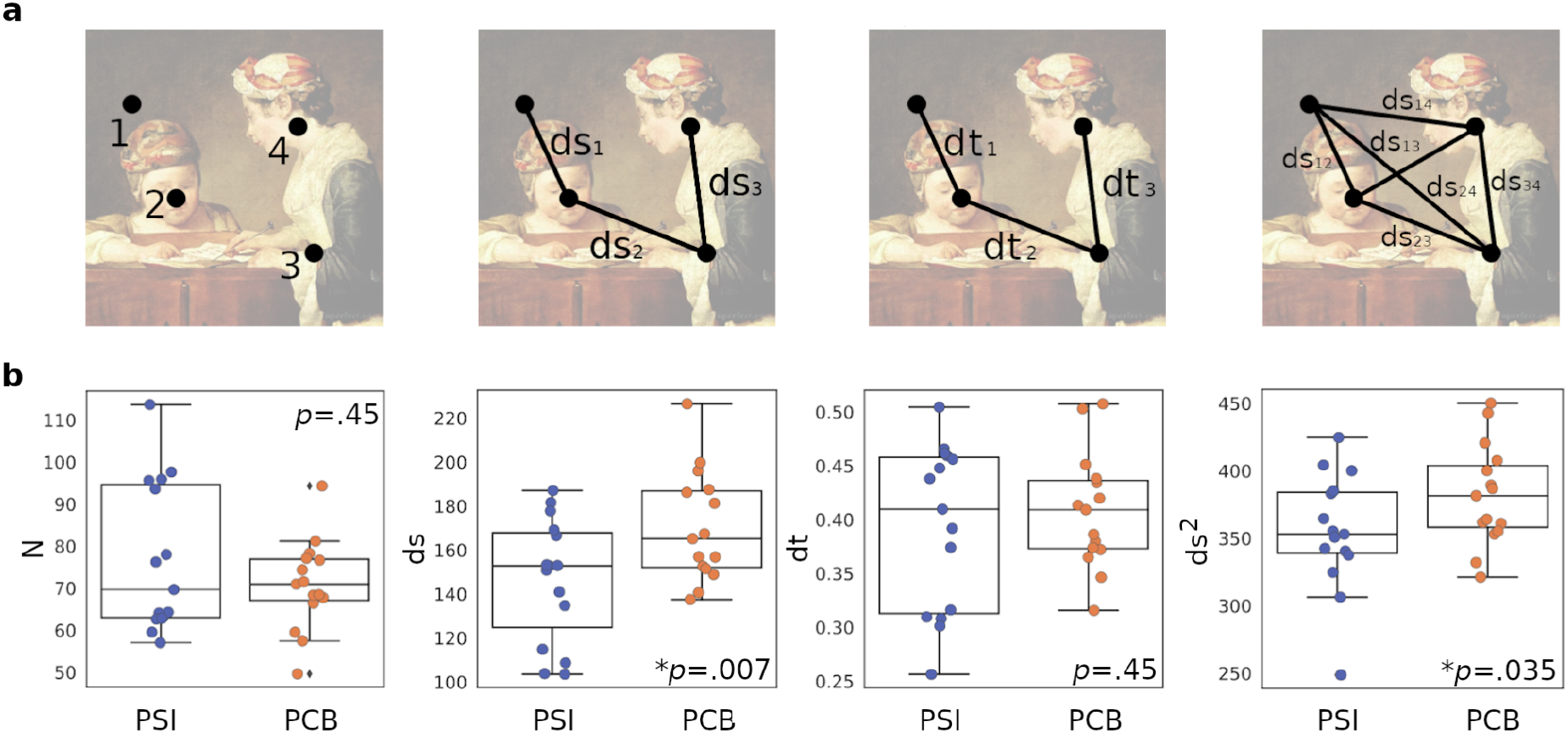
Changes in fixation metrics in high vs. low dose conditions. **a)** Illustration of the statistical fixation metrics that were calculated for each image. From left to right: number of fixations (N), distance between consecutive fixations (*ds*), time between consecutive fixations (*dt*) and distance between all pairs of fixations (*ds*^2^). **b)** Values of the metrics N, *ds, dt* and *ds*^2^, averaged across stimuli for each subject, and compared between the high dose of psilocybin (PSI) vs. the active placebo (PCB) condition. The p-values (Wilcoxon signed-rank tests) are shown as insets.

### Statistical fixation metrics

As described in the Methods, the total number of fixations (*N*), the average distance between fixations (*ds*), the average time between fixations (*dt*) and the average distance between all pairs of fixations (*ds*^2^) were calculated for each stimulus and averaged across all stimuli for each participant and dosing condition. The resulting values were compared between dosing conditions using non-parametric paired Wilcoxon signed-rank tests (Figure 4b). A significant effect of the dosing condition was found for the metrics *ds* and *ds*^2^, in both cases indicative of lower values for the high dose condition.

The standard deviation of the distances between consecutive fixations (*σ*(*ds*)) and the standard deviation of the distances between all pairs of fixations (*σ*(*ds*^2^)) were calculated and averaged across images for each participant and dosing condition. For both metrics, the statistical analysis also revealed significantly lower values for the high dose condition (Figure 5).

**Figure 5.**
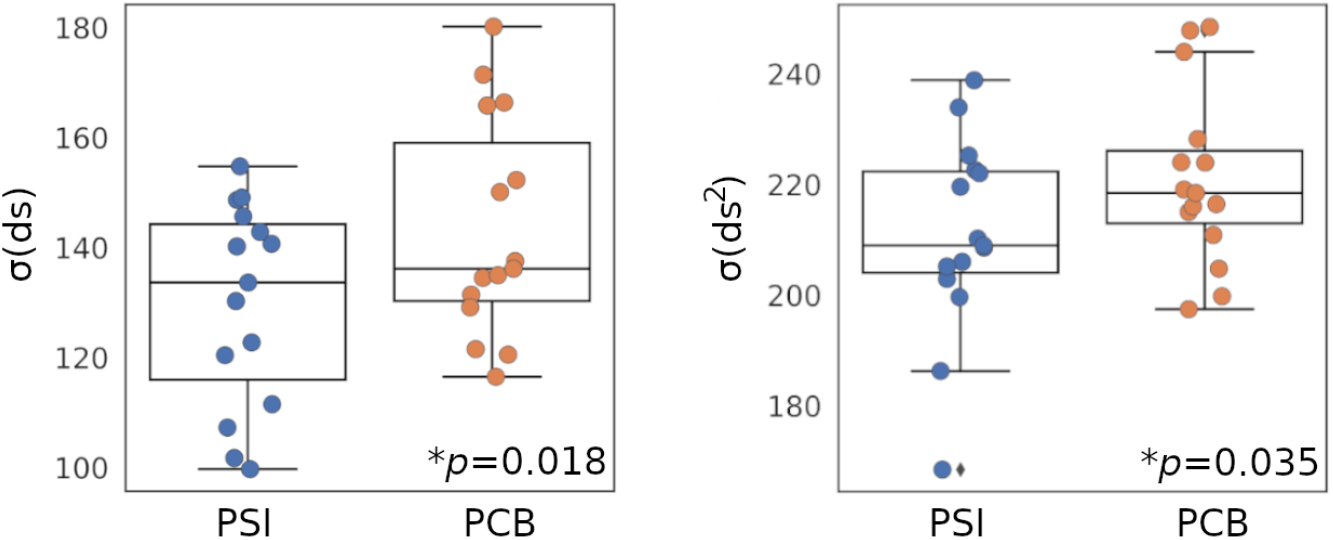
Average values of the standard deviations of the distances between consecutive fixations, *ds*, (left) and of the distances between all pairs of fixations, *ds*^2^ (right). Results are shown for the high dose of psilocybin (PSI) and active placebo (PCB) conditions, and compared using Wilcoxon signed-rank tests (p-values shown as insets).

### Shannon Entropy

Figure 6 shows the Shannon entropy calculated for both conditions using four different bin widths. For each panel, the resulting fixation probability distribution is shown on the left, while the boxplots with the comparison between high and low dose conditions is shown on the right. Significant results were obtained for all bin widths, indicating lower entropy values for the high dose condition.

**Figure 6.**
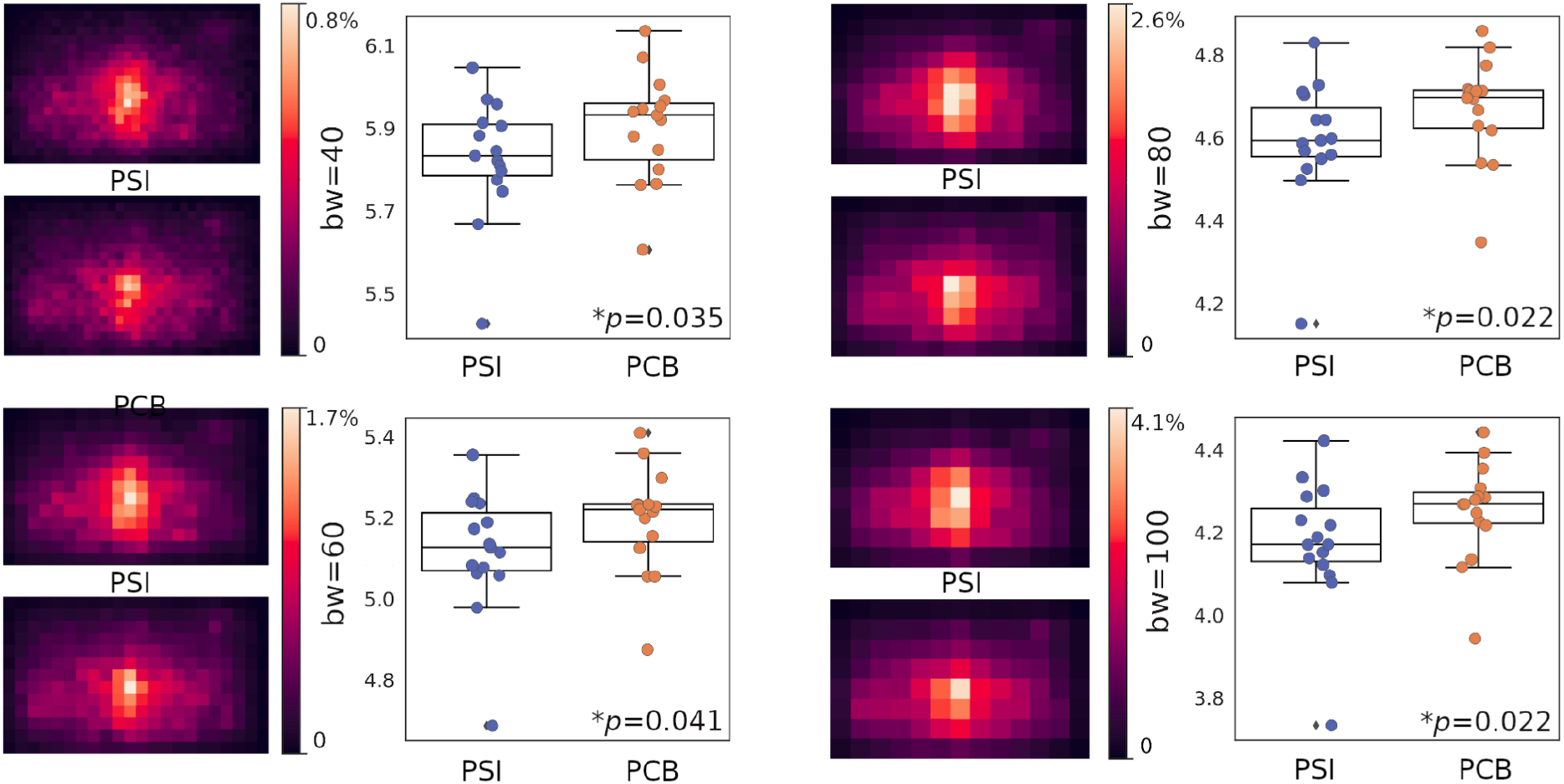
Psilocybin reduces the Shannon entropy of the fixation probability distribution. Panels show results obtained for the following bin width (bw) values: 40, 60, 80 and 100. For each panel, the resulting fixation probability distribution is shown on the left, while the boxplots with the comparison between high (PSI) and low dose (PCB) conditions is shown on the right. The p-values computed using Wilcoxon signed-rank tests are shown as insets.

Finally, we investigated whether correlations existed between the subjective reports (Figure 3) and the fixation metrics assessed using eye tracking (Figure 4). For this purpose, we computed the Pearson linear correlation coefficient between the metrics *N, ds, dt* and *ds*^2^ and the AEQ dimensions, both for the high and low dose conditions. We focused on cases where at least one of the conditions presented a correlation with high effect size (r≥0.5), which only occurred for the correlation between *ds* and the emotional response sub-scale of the AEQ. As shown in Figure 7, a significant correlation of r=0.71 was found for the low dose condition; however, no correlation between these variables existed during the high dose condition. Moreover, the difference between these correlation coefficients was significant (Hinkle et al. 2003).

**Figure 7.**
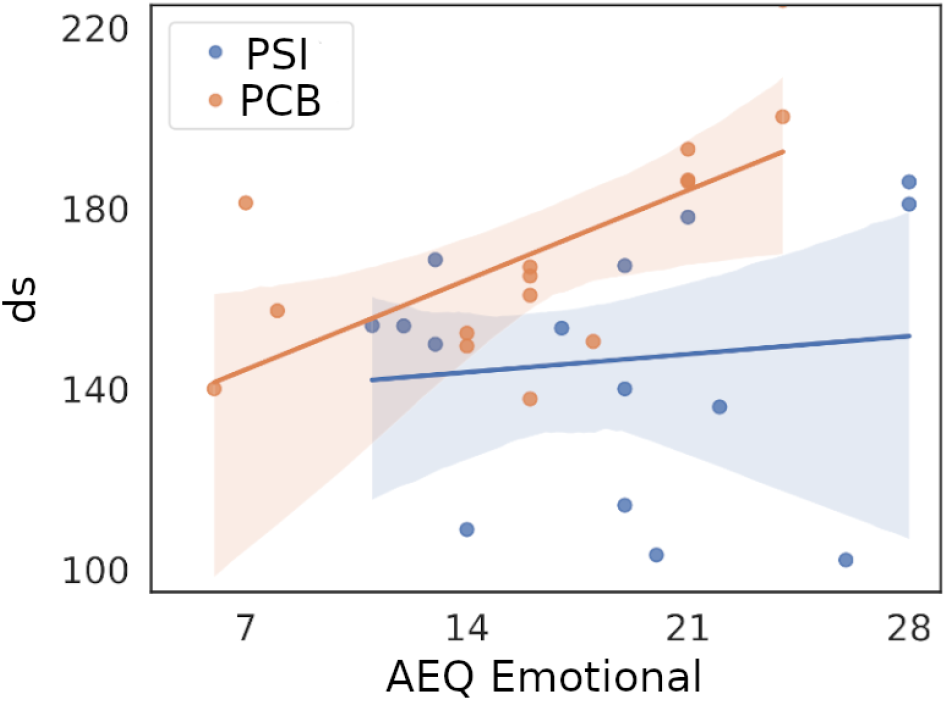
Scatter plot of the fixation metric vs. the emotional sub-scale of the AEQ, including the best linear fit in the least squares sense for the low (PCB) and high (PSI) dose conditions. The correlation coefficient obtained for the high dose condition was r=0.71, while the value for the low dose condition was r=0.079, resulting in a significant increase of the Pearson correlation from the low to the high dose condition.

## Discussion

We implemented double-blind placebo-controlled paradigm under naturalistic conditions to investigate the modulation of perception by psilocybin, a natural serotonergic psychedelic of major contemporary interest in basic and clinical neuroscience. Instead of focusing solely on the measurement and analysis of subjective reports, we used eye tracking to quantify the behavior of participants regarding the statistics of their visual fixations when presented paintings from different art periods. These stimuli are naturally engaging and diverge from everyday visual scenes, thus factoring in an element of uncertainty in the experiment; moreover, their use represents an opportunity to further our knowledge of how psychedelic compounds influence aesthetic perception, a long-standing field of research which has gained attention recently (van Elk et al., 2021; Aday et al., 2023). Our results suggest that participants under the high dose acquire visual information more locally, with shorter distances between all fixation pairs, as well as between consecutive fixations. Psilocybin also reduced the variability in these two metrics, while decreasing the entropy of the fixation probability distribution, which is consistent with narrower spatial distribution of attention under the high dose condition. Finally, we found that psilocybin induced a decorrelation between the consecutive fixation distance and the emotional response dimension of the aesthetic experience, as measured with the AEQ. With regards to the aesthetic experience itself, participants reported heightened emotional response and state of flow under the influence of the high dose relative to the control condition.

Ample anecdotal evidence suggests a link between the output of successful artists and the use of psilocybin, LSD and other serotonergic psychedelics, contributing to the emergence of the movement known as “psychedelic art” (Lee & Shlain, 1992). However, few rigorous studies attempted to clarify the relationship between the acute effects of psychedelics, artistic perception and expression, and creativity. A recent naturalistic study addressed the effects of psychedelic microdosing on aesthetic perception (van Elk et al. 2022), failing to find an effect of drug intake on feelings of awe elicited by paintings, thus partially contradicting anecdotal reports of a deeper and more profound aesthetic perception caused by these substances. Even though we did not explicitly address the construct of awe, we found increased scores of the emotional response dimension of the AEQ during the high dose condition. We also found a marginally significant effect of psilocybin on the flow of experience dimension. Interestingly, previous work suggests some degree of overlap between the constructs of flow and absorption (Marty-Dugas and Smilek 2019), the latter being originally defined as “a disposition for having episodes of total attention that fully engage one’s representational resources” (Tellegen and Atkinson 1974). It is known that trait absorption and the acute effects of psychedelics can exert mutual influences (Studerus et al. 2012; Haijen et al. 2018; Aday et al. 2021), and that absorption plays an important role in the processing of aesthetic stimuli from different modalities (Combs et al. 1988; Rhodes et al. 1988; Kuijpers et al. 2019). However, the lack of studies using eye tracking hinders a direct comparison between these studies and our findings.

Recent work from Aday and colleagues (Aday et al., 2023) investigated changes in aesthetic experiences elicited by ayahuasca, adopting an open label naturalistic design. Measured with the AEQ, participants reported increased levels of aesthetic experience in two follow-up sessions (one week and one month after the ayahuasca retreat). These changes occurred for all dimensions of the questionnaire, while in our study we only found differences for two facets. This discrepancy could be attributed to expectation effects in the study conducted by Aday et al., given the lack of a control condition, or due to the more intense and sustained nature of psychedelic use in that study, which consisted of retreats totaling between 2 and 7 ayahuasca sessions. Another factor to be considered is a possible effect of the setting on aesthetic perception, as natural environments could favor aesthetic experiences more than urban contexts. Finally, neither our experiment nor the one conducted by Aday and colleagues revealed significant correlations between acute effects (e.g. mystical type experiences) and the changes in the AEQ scores. Further research is required to understand how aesthetic perception relates to other idiosyncratic aspects of the psychedelic experience.

The reported changes in fixation metrics could stem from the effects of psilocybin on low-level visual perception, top-bottom attentional processes, or a combination of both. According to a model proposed by Chatterjee (2003) the first stage of aesthetic perception corresponds to the identification of lower-level variables (color, form, texture, etc.), which are subsequently integrated to form a holistic view, directing visual attention to specific areas of interest depending on visual composition and complexity. The well-established effects of psychedelics on low-level visual perception (Preller & Vollenweider, 2018) could influence the first stage of this process, while the acute effects on attention could influence the second stage (Bălăeţ 2022; van Elk & Yaden 2022). Previous studies established that more global fixation distributions are characteristic of the judgment of aesthetic appeal, while more local distributions relate to the processing of visual complexity in the painting (Wallraven et al. 2009), suggesting that our findings could stem from the interaction between psychedelics and the perception of low-level visual features in the stimuli. The positive correlation between *ds* and the AEQ sub-scale assessing the emotional aspect of the aesthetic experience suggests that semantic content is gathered by exploring the painting more globally when the participant is more emotionally engaged with the painting. Conversely, the decorrelation between these variables observed for the high dose condition could indicate that the emotional involvement with the artwork, which is overall greater in this condition, does not affect the way it is explored, suggesting that the increase in emotional engagement is not directly related to the semantic content gathered during visual exploration. However, it must be noted that the immediate VAS scales administered after each painting included a rating of subjective aesthetic appeal, yet no significant effects of psilocybin were found on this variable.

According to the REBUS theory proposed by Carhart-Harris and Friston, psychedelics act by relaxing the precision of high-level priors or beliefs, thereby liberating bottom-up information flow (Carhart-Harris and Friston 2019). In visual perception, this relaxation could manifest as a more entropic exploration of the visual scene, as the priors fail to inform gaze direction to regions where the majority of the relevant semantic information is represented. To date, this hypothesis has received evidence from multiple experiments, including complex tasks such as the production of natural speech (Carhart-Harris and Friston 2019; Sanz et al. 2021). However, it appears to contradict the main findings of our study. As mentioned above, this could be caused by the effects of psilocybin on the processing of low-level visual features, causing an exaggerated interest in this aspect of the paintings at the expense of a more global appreciation (Wallraven et al. 2009). In The Doors of Perception, Aldous Huxley famously narrated how his attention was lost at seemingly insignificant details of the visual scene: “I looked down by chance, and went on passionately staring by choice, at my own crossed legs. Those folds in the trousers—what a labyrinth of endlessly significant complexity! And the texture of the grey flannel—how rich, how deeply, mysteriously sumptuous!” (Huxley 1956). Such mode of perception seems compatible with the more local exploration revealed by our analysis of the eye tracking data.

The interpretation of our results faces some limitations, mainly due to specific choices concerning the experimental design. In this study, we prioritized the inclusion of tasks which are either natural and/or spontaneous (e.g. production of language), or capable of engaging the participants for relatively long periods of time, as is the case for the visual perception and the aesthetic appreciation of artworks. We also opted to pursue this experiment under conditions that are less constrained and more natural than those of a typical lab-based study. The merits and drawbacks of this approach have been discussed with detail elsewhere (Haijen et al. 2018; van Elk et al. 2022; Sanz et al. 2022; Tagliazucchi 2022), yet here it is important to highlight that distractions during an eye tracking task can easily introduce a large amount of missing data points, potentially rendering the gathered data useless for further analysis. This issue could be expected to become more problematic in settings where subjects can develop feelings of fear and/or anxiety, including research laboratories and hospital settings (Studerus et al. 2012). Another characteristic of our study is the inclusion of a large variety of paintings, as opposed to presenting a reduced repertoire for a longer duration. In particular, this choice could have limited the processing of the stimuli to that of low-level features, as discussed above. We decided to present paintings for comparatively short durations to maintain the attention and engagement of the participants. Another potential limitation could arise due to the large unblinding rate of the conditions, compatible with previous studies (van Elk et al. 2022) even when adopting an active condition as control (i.e. the low dose). However, the nature of the eye tracking recordings themselves make it unlikely that participants could bias their responses based on their perception of the experimental condition, as they are objective metrics that capture complex patterns of eye movements. Finally, a major limitation of our study is its exploratory nature. In this regard, we consider that exploratory studies are necessary when previous literature is scarce - in this particular case, without reports addressing the sensitivity of eye tracking to capture behavioral changes elicited by psychedelics (Jaeger and Halliday 1998). Future work by our group and others should build upon this first effort and narrow the conclusions by studying more specific aspects of visual perception.

In conclusion, we achieved a quantitative description of fixation statistics during aesthetic perception and their modulation by mushrooms of the *Psilocybe* genus, while also shedding light on the interaction between acute psychedelic effects and the aesthetic experience elicited by a selection of paintings. Guided by theoretical efforts positing a relaxation of priors during the acute effects of serotonergic psychedelics, we first hypothesized a more entropic distribution of fixations. However, we found the opposite result, which could be compatible with the idiosyncratic effects of psilocybin on the perception of low-level visual information. Future studies are required to address this and other possibilities raised by our work, a first attempt to quantify the acute effects of psychedelics on the complex eye movement behaviors underlying visual perception.

## Acknowledgments

The authors acknowledge funding received from Spinoza B.V. to support this project.

## Supplementary material for

### 1. Self-reported scales and questionnaires

Big Five Inventory (BFI). A validated Spanish version of the inventory assessing five dimensions of personality: neuroticism, extraversion, openness to experience, agreeableness, and conscientiousness (Benet-Martínez and John 1998). The BFI questionnaire consists of 44 items based on a 5-point Likert scale. Multiple studies suggest that psychedelics are capable of inducing short and long-term changes in personality (Bouso et al. 2018). Moreover, some of these changes (e.g. increased openness) might be contributing factors to the therapeutic effects of psychedelics, as well as to the long-term positive changes in subjective well-being reported by healthy Individuals

Short Suggestibility Scale (SSS). An inventory that assesses suggestibility, created by Kotov et al (2004), and translated to Spanish by the authors. The questionnaire consists of 21 items based on a 5-point Likert scale. It has been shown that psychedelics can enhance suggestibility in healthy volunteers (Carhart-Harris et al. 2015)

Tellegen Absorption Scale (TAS). A 34-item scale developed to measure the capacity of an individual to become absorbed in the performance of a task (Tellegen and Atkinson 1974), translated to Spanish by the authors. This psychological construct is positively correlated with the overall intensity of the effects elicited by psychedelic drugs, and also implicated in some of its most intriguing effects, such as the induction of spiritual or mystical experiences (Haijen et al. 2018)

State-Trait Anxiety Inventory (STAI-T / STAI-S). Validated Spanish editions of commonly used scales which measure state anxiety (situational anxiety of a temporary nature) and trait anxiety (stable trait linked to individual characteristics) (Spielberger 1983). The instrument comprises 40 items and is based on a 4-point Likert scale.

Positive and Negative Affect Schedule (PANAS). A validated Spanish version of a psychometric scale that has been widely used to measure dimensions of affect, both positive and negative (Watson et al. 1988). The instrument consists of 20 affirmations based on a 5-point Likert scale.

Psychological Well-being Scale (BIEPS). A scale used to measure eudaemonic well-being in adults (including dimensions of acceptance, perception of control, social ties, and autonomy and projects) (Castro 2002) It consists of 13 questions based on a 3-point Likert scale.Originally developed in Spanish.

Pre-ceremony Scale (PRE). Twelve items assessing non-pharmacological contextual factors prior to psychedelic experiences. Using principal component analysis, Haijen et al. showed that the items clustered into three components: set, setting and clear intentions.(Haijen et al. 2018).

Expectation (EXP). Seventeen questions designed to measure expectations of change in the following areas: positive emotions, negative emotions, anxiety, attention, absorption, creativity, perception, problem solving, empathy, memory, energy, sleep, sociability, spirituality, openness, oceanic feeling and substance intake (Cavanna et al. 2022).

Altered States of Consciousness (5D-ASC). Ninety-four items assessing different aspects of altered states of consciousness, understood as temporary deviations from normal waking consciousness. Consists of three main dimensions, each with several lower-order scales. The dimensions and their corresponding scales are: oceanic boundlessness (experience of oneness, spiritual experience, state of bliss, insightfulness), anxious ego dissolution (disembodiment, impaired control and cognition, anxiety), visionary restructuration (complex images, elementary images, audiovisual synesthesia, altered meaning of perceptions) (Studerus et al. 2010).

Mystical Experiences Questionnaire (MEQ-30). Thirty items from which four subscale scores are calculated: mystical, positive mood, transcendence of time and space, and ineffability, which are considered the most relevant and defining aspects of mystical experiences (Barrett et al. 2015).

Aesthetic Experience Questionnaire (AEQ). Four art-related factors - Perceptual, Emotional, Cultural and Understanding - and two flow factors - Proximal Conditions and Experience - make up this questionnaire, which assesses aspects of aesthetic experience in a general population (Wanzer et al. 2018). Emotional factor: assesses the individual’s range of emotions, how they change over time, how they are moved, and how they experience physical reactions when viewing art. Cultural factor: examines the individual’s engagement with the historical and cultural context of the artwork, perceiving it as representative of its time, placing it in a historical context and relating it to other artworks. Perceptual factor: assesses the individual’s attention to the composition and colors of the artwork, as well as their focus on the subtle aspects and details of the artwork. Understanding factor: measures the individual’s effort to fully comprehend and interpret the artwork, to understand the artist’s intended message, to gain new insights specifically about the artwork, and to perceive the artwork as an extension of the artist. Flow-Proximal Conditions factor: examines the clarity of the individual’s goals in viewing the artwork, their confidence in their interpretations, and their sense of being able to understand and engage with the artwork. Flow-Experience factor: assesses the individual’s experience of losing track of time, being absorbed in thought, maintaining focus and concentration, and finding the experience of viewing the artwork rewarding and enjoyable.

Emotional valence and beauty rating VAS. Self reported scales assessing the individual’s valuation of a painting. For the emotional valence VAS, the following statement is presented: “The emotion I felt about this painting was…”, with 7 possible responses ranging from: “Very negative” to “Very positive” . For the beauty rating VAS, the following statement is presented: “This painting is very beautiful… “, with 7 possible responses ranging from “I strongly disagree” to “I strongly agree.”

### 2. Summary of statistical analyses

The results of all the self-reported scales and questionnaires are presented in table S1. The results of the measurements during the acute effects are presented in table S2.

**Table S1.** Baseline, Pre dose, Acute effects and Post dose outcomes are presented as mean ± standard deviation. Statistical significance is indicated using asterisks and computed using Student’s t-test.

| Scale - Factor | Mean ± SD<br>Psilocybin | Mean ± SD<br>Placebo | p-value | Cohen's d | BF<sub>10</sub> |
| --- | --- | --- | --- | --- | --- |
| **Baseline** |  |  |  |  |  |
| STAI - Trait | 21.04 ± 8.39 | 20.61 ± 8.04 | 0.645 | 0.05 | 0.24 |
| BFI - Extraversion | 28.61 ± 4.15 | 28.57 ± 3.80 | 0.945 | 0.01 | 0.22 |
| BFI - Agreeability | 34.26 ± 4.85 | 34.48 ± 4.87 | 0.623 | 0.04 | 0.25 |
| BFI - Conscientiousness | 30.96 ± 5.47 | 30.00 ± 6.54 | 0.28 | 0.16 | 0.38 |
| BFI - Neuroticism | 21.78 ± 7.39 | 20.87 ± 6.36 | 0.166 | 0.13 | 0.54 |
| BFI - Openness | 43.48 ± 4.59 | 42.83 ± 5.58 | 0.303 | 0.13 | 0.36 |
| TAS | 21.83 ± 6.14 | 20.57 ± 6.73 | 0.158 | 0.2 | 0.56 |
| SSS | 56.48 ± 11.24 | 53.26 ± 10.21 | 0.077 | 0.3 | 0.94 |
| **Pre dose** |  |  |  |  |  |
| STAI - State | 11.26 ± 8.62 | 12.09 ± 8.38 | 0.692 | 0.1 | 0.23 |
| PANAS - Negative affect | 15.32 ± 5.16 | 14.09 ± 4.46 | 0.274 | 0.26 | 0.38 |
| PANAS - Positive affect | 32.57 ± 9.31 | 34.30 ± 7.86 | 0.268 | 0.2 | 0.39 |
| BIEPS | 35.48 ± 2.87 | 35.55 ± 2.86 | 0.875 | 0.02 | 0.22 |
| PRE - Set | 6.63 ± 0.66 | 6.60 ± 0.79 | 0.861 | 0.05 | 0.22 |
| PRE - Setting | 8.54 ± 1.10 | 8.49 ± 1.50 | 0.816 | 0.03 | 0.22 |
| PRE - Clear intentions | 5.74 ± 1.96 | 5.67 ± 2.87 | 0.839 | 0.03 | 0.22 |
| EXP - Negative emotions | 2.00 ± 0.80 | 2.00 ± 0.80 | 1.000 | 0 | 0.22 |
| EXP - Anxiety | 1.91 ± 0.90 | 2.04 ± 1.02 | 0.623 | 0.14 | 0.25 |
| EXP - Attention | 3.13 ± 1.32 | 3.30 ± 1.18 | 0.505 | 0.14 | 0.27 |
| **EXP - Absorption** | **4.17 ± 0.72** | **3.74 ± 0.54** | **0.009 \*\*** | **0.68** | **5.36** |
| EXP - Creativity | 4.17 ± 0.58 | 3.96 ± 0.77 | 0.171 | 0.32 | 0.53 |
| EXP - Perception | 4.50 ± 0.66 | 4.26 ± 0.62 | 0.102 | 0.37 | 0.77 |
| EXP - Problem resolution | 3.26 ± 1.39 | 3.35 ± 1.07 | 0.704 | 0.07 | 0.23 |
| EXP - Empathy | 4.39 ± 0.58 | 4.13 ± 0.81 | 0.056 | 0.37 | 1.21 |
| EXP - Memory | 2.91 ± 1.12 | 3.09 ± 0.95 | 0.406 | 0.17 | 0.30 |
| EXP - Energy | 3.74 ± 1.10 | 3.61 ± 0.89 | 0.575 | 0.13 | 0.25 |
| EXP - Sleep | 2.74 ± 1.01 | 2.61 ± 0.89 | 0.503 | 0.14 | 0.27 |
| **EXP - Sociability** | **3.83 ± 0.94** | **3.35 ± 1.07** | **0.038 (\*)** | **0.48** | **1.64** |
| EXP - Spirituality | 4.14 ± 0.69 | 3.96 ± 0.71 | 0.204 | 0.26 | 0.47 |
| EXP - Openness | 4.39 ± 0.78 | 4.13 ± 0.69 | 0.162 | 0.35 | 0.55 |
| **EXP - Oceanic feelings** | **4.61 ± 0.58** | **4.17 ± 0.72** | **5e-4 (\*\*\*)** | **0.67** | **70.52** |
| EXP - Substance intake | 2.65 ± 0.83 | 2.83 ± 0.78 | 0.406 | 0.22 | 0.30 |
| EXP - Total | 60.57 ± 5.69 | 58.61 ± 6.46 | 0.108 | 0.32 | 0.73 |
| **Acute effects** |  |  |  |  |  |
| **AEQ - Emotional** | **18.38 ± 5.55** | **15.26 ± 5.79** | **0.004 (\*\*)** | **0.55** | **11.15** |
| AEQ - Cultural | 19.48 ± 6.12 | 20.74 ± 6.35 | 0.341 | 0.2 | 0.33 |
| AEQ - Perceptual | 16.14 ± 3.53 | 16.78 ± 4.08 | 0.494 | 0.17 | 0.27 |
| AEQ - Understanding | 17.38 ± 5.36 | 18.04 ± 4.81 | 0.658 | 0.13 | 0.24 |
| AEQ - Flow conditions | 11.19 ± 4.38 | 10.22 ± 2.80 | 0.287 | 0.27 | 0.37 |
| **AEQ - Flow experience** | **19.81 ± 4.37** | **16.78 ± 4.65** | **0.015 (\*)** | **0.67** | **3.55** |
| **Post dose** |  |  |  |  |  |
| STAI - State | 10.79 ± 7.97 | 8.43 ± 7.25 | 0.128 | 0.31 | 0.65 |
| **PANAS - Negative affect** | **16.23 ± 5.43** | **14.26 ± 5.82** | **0.108** | **0.35** | **0.73** |
| PANAS - Positive affect | 32.68 ± 9.94 | 34.61 ± 7.77 | 0.277 | 0.22 | 0.38 |
| BIEPS | 8.41 ± 0.89 | 8.57 ± 0.73 | 0.314 | 0.19 | 0.35 |
| MEQ - Mystical experience | 21.38 ± 15.48 | 4.48 ± 6.49 | 4e-6 (***) | 1.42 | 4565 |
| MEQ - Positive mood | 14.41 ± 6.34 | 6.52 ± 4.39 | 4e-5 (***) | 1.45 | 686 |
| MEQ - Transcendence | 12.45 ± 5.44 | 2.35 ± 2.57 | 2e-7 (***) | 2.38 | 91510 |
| MEQ - Ineffability | 8.23 ± 3.67 | 1.57 ± 1.70 | 3e-8 (***) | 2.33 | 435700 |
| 5DASC - Unity | 14.83 ± 11.28 | 6.35 ± 1.97 | 0.001 (**) | 1.05 | 31.07 |
| 5DASC - Spiritual | 8.13 ± 5.29 | 4.13 ± 2.34 | 0.003 (**) | 0.98 | 15.71 |
| 5DASC - Blissful | 12.23 ± 8.18 | 5.39 ± 3.14 | 0.001 (***) | 1.1 | 53.24 |
| 5DASC - Insightfulness | 12.59 ± 8.34 | 4.91 ± 3.80 | 6e-5 (***) | 1.18 | 454 |
| 5DASC - Disembodiment | 7.27 ± 5.55 | 3.57 ± 1.50 | 0.003 (**) | 0.91 | 13.77 |
| 5DASC - Impaired cognition | 22.86 ± 10.48 | 8.57 ± 3.01 | 1e-6 (***) | 1.86 | 15920 |
| 5DASC - Anxiety | 17.50 ± 9.70 | 7.26 ± 3.02 | 9e-5 (***) | 1.42 | 294 |
| 5DASC - Complex imagery | 15.73 ± 9.10 | 4.30 ± 2.98 | 2e-6 (***) | 1.69 | 8747 |
| 5DASC - Elementary imagery | 20.27 ± 10.22 | 4.57 ± 3.86 | 6e-7 (***) | 2.03 | 30910 |
| 5DASC - Audio-visual syn. | 13.41 ± 11.65 | 4.65 ± 3.95 | 0.002 (**) | 1.01 | 24.12 |
| 5DASC - Changed meaning | 14.36 ± 7.85 | 4.13 ± 2.20 | 1e-6 (***) | 1.78 | 13850 |

**Table S2.** Acute effects outcomes are presented as mean ± standard deviation.

| Scale - Factor | Mean ± SD<br>Psilocybin | Mean ± SD<br>Placebo | p-value | RBC's r |
| --- | --- | --- | --- | --- |
| **Acute Effects** |  |  |  |  |
| VAS 1 total | 17.56 ± 8.47 | 9.91 ± 3.80 | 0.002 ** | 0.80 |
| VAS 2 total | 21.83 ± 7.79 | 11.87 ± 5.19 | 0.0001*** | 0.84 |
| VAS 3 total | 20.00 ± 7.56 | 9.35 ± 2.76 | 8e-5 *** | 0.96 |
| VAS 4 total | 14.95 ± 8.00 | 7.91 ± 1.35 | 0.001 ** | 0.87 |
| Emotional valence | 0.52 ± 0.35 | 0.41 ± 0.35 | 0.232 | 0.30 |
| Beauty rating | 1.15 ± 0.6 | 1.13 ± 0.51 | 0.708 | 0.10 |
| N | 77.00 ± 17.19 | 70.80 ± 10.32 | 0.454 | 0.23 |
| ds | 146.43 ± 27.42 | 170.19 ± 23.89 | 0.007** | -0.76 |
| dt | 0.39±0.08 | 0.41±0.05 | 0.454 | -0.23 |
| ds2 | 354.78±41.93 | 381.87±36.33 | 0.035* | -0.62 |
| σ(ds) | 129.93±17.68 | 143.22±19.32 | 0.018* | -0.68 |
| σ(ds2) | 210.56±17.36 | 220.96±15.35 | 0.035* | -0.62 |
| Shannon entropy (bw=40) | 5.83±0.14 | 5.90±0.13 | 0.035* | -0.62 |
| Shannon entropy (bw=60) | 5.12±0.15 | 5.19±0.13 | 0.041* | -0.60 |
| Shannon entropy (bw=80) | 4.59±0.15 | 4.67±0.12 | 0.022* | -0.67 |
| Shannon entropy (bw=100) | 4.18±0.15 | 4.25±0.12 | 0.022* | -0.67 |

### 3. Stimuli

The following is a list of the authors, titles and completion years of the artworks displayed in the eye tracking task.

1. Jan van Eyck: The Madonna of the Chancellor Rolin (1434)
2. Hugo van der Goes: Adoration of the Kings (around 1470)
3. Sandro Botticelli: The Birth of Venus (1478–1487)
4. Albrecht Dürer: Self-portrait (1498)
5. Pieter Brueghel the Elder: Landscape with the Fall of Icarus’ (c. 1550)
6. Jan Matsys: Flora (1559)
7. El Greco: View of Toledo (1600–1610)
8. Hendrick Avercamp: Winter Scene on a Canal (c. 1630)
9. Francisco de Zurbarán: Still Life: Lemons, Oranges and a Rose (1633)
10. Hyacinthe Rigaud: Portrait of Louis XIV. (1701)
11. Jean Siméon Chardin: The Young Schoolmistress (before 1740)
12. Françis Boucher: Marie-Louise O’Murphey (1751)
13. Caspar David Friedrich: Polar Sea (1822–1824)
14. Carl Spitzweg: The Poor Poet (1839)
15. William Holman Hunt: The Hireling Shepherd (1851)
16. Thomas Eakins: Max Schmitt in a Single Scull (1871)
17. Gustave Caillebotte: Parisian Street, Rainy Day (1877)
18. Vincent van Gogh: Café Terrace at Night (1888)
19. Paul Cézanne: La Montagne Sainte-Victoire (1897)
20. Winslow Homer: The Fox Hunt (1893)
21. Ferdinand Hodler: Youth Admired by the Woman (1903)
22. Pierre Bonnard: Backlit Nude (1908)
23. Marcel Duchamp: Sad Young Man in a Train (1911)
24. Giacomo Balla: Abstract Speed + Sound (1913–14)
25. Grant Wood: American Gothic (1930)
26. Otto Dix: Flanders (1934–1936)
27. Pablo Picasso: Guernica (1937)
28. Paul Nash: Dream Landscape (1936–1938)
29. Francis Bacon: Three Studies for Figures at the Base of a Crucifixion (1944)
30. Helen Frankenthaler: Mountains and Sea (1952)

